# Social-ecological factors and preventive actions decrease the risk of dengue infection at the household-level: results from a prospective dengue surveillance study in Machala, Ecuador

**DOI:** 10.1101/136382

**Authors:** Aileen Kenneson, Efraín Beltrán-Ayala, Mercy J. Borbor-Cordova, Mark E. Polhemus, Sadie J. Ryan, Timothy P. Endy, Anna M. Stewart-Ibarra

## Abstract

**Background:** In Ecuador, dengue virus (DENV) infections transmitted by the *Aedes aegypti* mosquito are among the greatest public health concerns in urban coastal communities. Community- and household-level vector control is the principal means of controlling disease outbreaks. This study aimed to assess the impact of knowledge, attitudes, and practices (KAPs) and social-ecological factors on the presence or absence of DENV infections in the household..

**Methods:** In 2014 and 2015, individuals with DENV infections from sentinel clinics were invited to participate in the study, as well as members of their household and members of four neighboring households located within 200 meters. We conducted diagnostic testing for DENV on all study participants; we surveyed heads of households (HOHs) regarding demographics, housing conditions and KAPs. We compared KAPs and social-ecological factors between households with (n=139) versus without (n=80) DENV infections, using bivariate analyses and multivariate logistic regression models with and without interactions.

**Results:** Significant risk factors in multivariate models included proximity to abandoned properties, interruptions in piped water, and shaded patios (p<0.05). Significant protective factors included use of mosquito bed nets, fumigation inside the home, piped water inside the home (p<0.05). In bivariate analyses (but not multivariate modeling), DENV infections was positively associated with HOHs who were male, employed, and of younger age than households without infections (p<0.05). DENV infections were not associated with knowledgeattitude, or reported barriers to prevention activities.

**Discussion:** Specific actions that can be considered to decrease the risk of DENV infections in the household include targeting vector control in highly shaded properties, fumigating inside the home, and use of mosquito bed nets. Community-level interventions include clean-up of abandoned properties, daily trash pick-up, and reliable piped water inside houses. These findings can inform interventions to reduce the risk of other diseases transmitted by the *Ae. aegypti* mosquito, such as chikungunya and Zika fever.

**Author summary:** Dengue, chikungunya and Zika viruses are transmitted to people primarily by the *Aedes aegypti* mosquitoes in tropical and subtropical regions. Diseases transmitted by the *Ae. aegypti* mosquito are a growing public health concern. Mosquito control is the principal means of preventing and controlling disease outbreaks. In this study, we compared the characteristics of households with and without DENV infections in the city of Machala, Ecuador. We found that risk factors for DENV infection included proximity to abandoned properties, interruptions in the piped water supply, and a highly shaded patio. Protective factors included the use of mosquito bed nets, fumigation inside the home, and piped water inside the home. These findings can be used to inform targeted vector control interventions by the public health sector at the household and community levels.

## INTRODUCTION

Dengue fever is a febrile illness caused by the *Flavivirus* dengue virus (DENV), of which there are four serotypes (DENV1-4) [1]. Infections may be asymptomatic, or have symptoms ranging from fever, rash, and joint pain to hemorrhage, shock and sometimes death. About 3.9 billion people, in 128 countries, are at risk of exposure to DENV infections [2,3]. In coastal Ecuador, the focus of this study, DENV infections and other febrile diseases transmitted by the *Aedes aegypti* mosquito, are among the greatest public health concerns. Over a five-year period (2010 to 2014), 72,060 cases of dengue illness were reported in Ecuador, with the highest incidence of cases in coastal urban areas [4]

*Aedes aegypti* is a tropical mosquito that has adapted to live and breed in urban environments [5,6]. *Ae. aegypti* also transmit chikungunya and Zika viruses, which now co-circulate with DENV in populations in the tropics and subtropics [7,8]. The female mosquitoes oviposit in water-bearing containers, which become the habitat of juvenile mosquitoes, such as water storage drums, tires, discarded containers, and flower pots [9–12]. Community- and household-level vector control interventions remain the principal means of controlling *Ae. aegypti*-borne disease outbreaks [13,14]. Preventive practices include covering water storage containers, eliminating standing water, adding larvicides to water containers, and general cleanup of potential water receptacles [1,14]. Placing screens on windows to protect against the mosquito vector has also been shown to be effective in preventing dengue transmission [15]. Indoor residential spraying in households has been shown to decrease the abundance of adult *Ae. aegypti* [16], and may decrease the risk of exposure to infected mosquitoes in households with DENV infections [17]. Novel vector control methods include lethal ovitraps [18], insect growth regulators (e.g., pyroproxyfen) [19], Wolbachia infections [20], and introduction of genetically-modified sterile mosquitoes into the population [21].

In Ecuador, the Ministry of Health (MOH) is the institution responsible for arbovirus and vector surveillance and control. Disease surveillance includes mandatory reporting of suspected (clinically diagnosed) and laboratory confirmed DENV cases. The vector control unit of the MOH is informed of new DENV cases from MOH clinics on average 8 days post diagnosis (range: 1 to 14 days) (*pers. comm.* T. Ordoñez). Focal vector control, including indoor residual spraying, is conducted in and around the households and neighboring households of people with DENV infections. Other regular vector control activities include a schedule of indoor residual spraying with deltamethrin and ultra-low volume with malathion fogging in high-risk urban communities at the beginning of the rainy season, and household visits by inspectors to treat water-bearing containers with an organophosphate larvicide (abate/temefos). Community cleanups occur before the rainy season to remove rubbish from household patios. Educational interventions for dengue prevention include television and radio campaigns, fliers, outreach to patients in MOH clinics, and community education meetings.

To improve the effectiveness of vector control and disease prevention interventions, public health practitioners require knowledge of local risk factors for dengue transmission. Early formative qualitative studies postulated that DENV infections were the result of underlying social structural inequities in urban areas, and they documented widespread misconceptions about dengue transmission and illness [22–24]. Similarly, in Ecuador, community members described dengue risk as the result of complex interactions among biophysical, political-institutional and community-household factors, such as climate conditions, low risk perception, economic barriers to prevention, lack of social cohesion, a lack of access to municipal services (e.g., piped water, sewerage, garbage collection), and failed coordination between municipal and public health authorities [25]. These and other studies indicated the need to frame dengue prevention in the context of broader social development goals through participatory multisectoral processes. Such efforts have proven to be complex (27) and require extensive community engagement (28).

To guide these interventions, studies were developed to assess dengue-related knowledge, attitudes, and practices (KAPs) and social-ecological risk factors. Dengue-related KAPs have been shown to be associated with the following demographic variables: sex, age, marital status, education, literacy, employment, occupation, income, ethnicity, and religion [26–37]. Other risk factors include frequent fogging of the neighborhood [38], adequate resources and assistance from public health staff [27], community support or governmental infrastructure to control neighboring and public spaces [39], and having a reputable source of information such as health personnel or head of the village [40]. In Ecuador, dengue risk was found to be associated with older female heads of household, access to piped water in the home, poor housing condition, household water storage, higher housing density, lack of knowledge, and low risk perception [41,42]. One of the limitations of KAP studies is that they often focus on preventative practices as the outcome of interest, rather than DENV infections. Other studies, such as those in Ecuador, utilize proxy variables for dengue risk (e.g., vector densities or MOH case reports), rather than DENV infections.

Active surveillance studies that detect symptomatic and subclinical DENV infections in the community provide a much more robust measure of DENV risk, especially when paired with direct observations of risk factors. Examples of enhanced surveillance studies that evaluated risk factors for DENV infections include a recent case-control study that was conducted in China [43], and seroprevalence studies that measured the presence of dengue antibodies (IgG, indicative of a past infection) in Malaysia, India, the Texas-Mexico border, Sudan, and Key West, Florida [44–49].

This study contributes to a growing body of knowledge on social-ecological and KAP risk factors for DENV infections. We detected acute and recent DENV infections through a passive and active surveillance study in Machala, Ecuador, in 2014 and 2015 [50]. We invited individuals with acute DENV infections (index cases) to participate in the study, along with other members of the index case household and members of four neighboring households located within 200 meters of the index house. We conducted diagnostic testing for DENV on all study participants, and we surveyed heads of households regarding household demographics, housing conditions, and dengue-related KAPs. Here, we present the results of analysis of the association between these risk factors and the presence or absence of DENV infections in households.

## METHODS

### Ethics Statement

This protocol was reviewed and approved by Institutional Review Boards (IRBs) at SUNY Upstate Medical University, the Luis Vernaza Hospital in Guayaquil, Ecuador, and the Ecuadorean MOH. Prior to the start of the study, all participants engaged in a written informed consent or assent process, as applicable. In the event the participant was unable to participate in the informed consent or assent process, a recognized health-care proxy represented them in the process and documented consent. The study population included children (>6 months) to adults.

### Study Site

Machala, Ecuador (population of approx. 276,000) is the capital city of El Oro Province and is a major port in the coastal lowland region (Figure 1). It is located 70 kilometers north of the Peruvian border. Like many cities in Latin America, people in the urban periphery have inadequate access to infrastructure and services, such as piped water and garbage collection, increasing their risk of DENV infections [41,42]. In 2014 and 2015, 1006 and 2877 suspected and confirmed DENV cases, respectively, were reported by the MOH, resulting in an incidence of 36.4 and 104 DENV cases per 10,000 people in the city of Machala. In 2014 all four DENV serotypes co-circulated, and in 2015 DENV1 and DENV2 were detected, along with the first cases of chikungunya. It should be noted that a high proportion of the suspected DENV cases in 2015 were misdiagnosed, and were actually chikungunya. Based on active surveillance in 2014 and 2015 [50], the prevalence of DENV infections is greatest in children and young adults under the age of 20, who accounted for 51% of all acute or recent DENV infections. For every medically-attended case, there are approximately three additional unreported DENV infections in the community.

**Figure 1.**
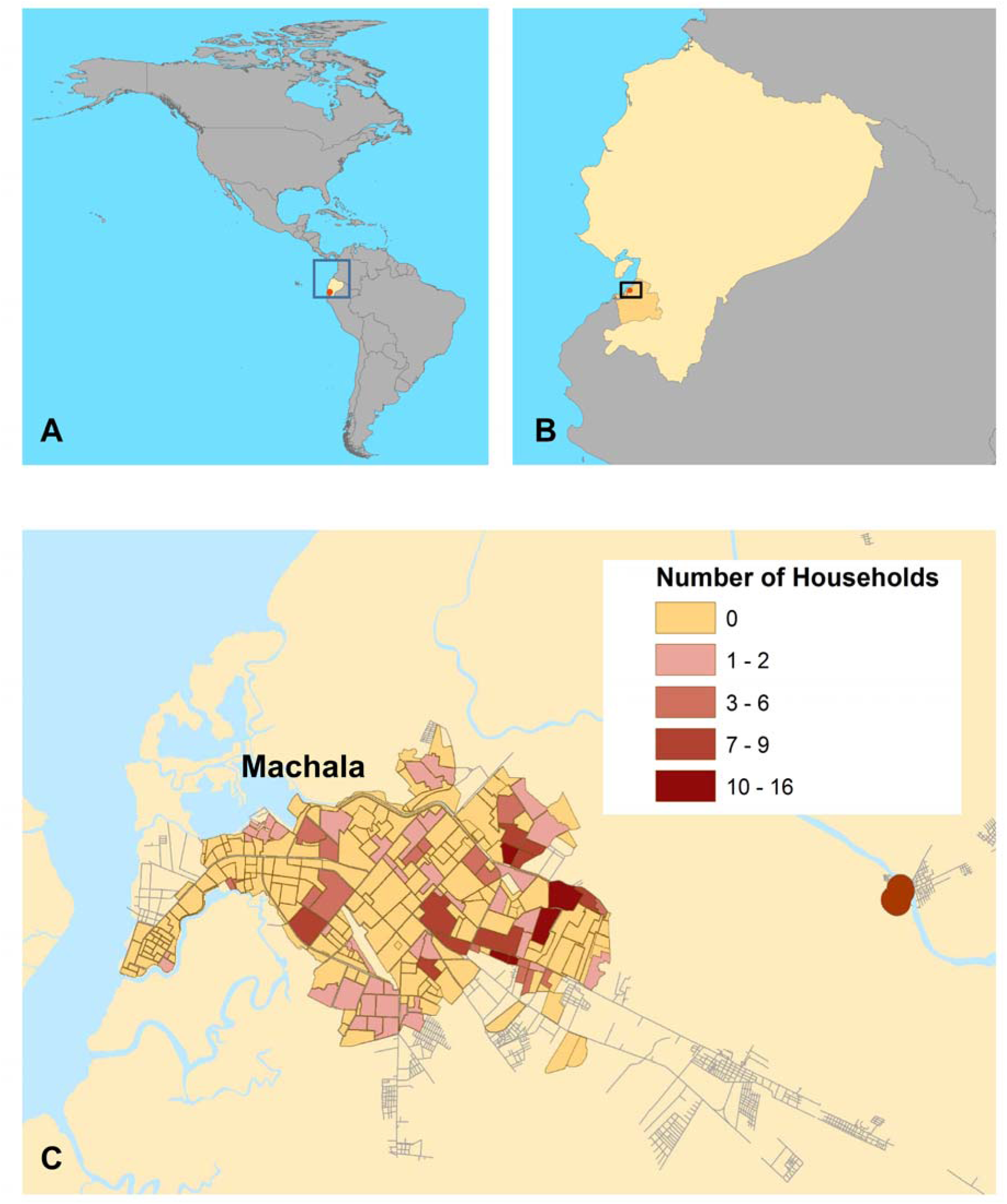
A map of the study site and distribution of study households. (A) location of Ecuador in the Americas (B) location of the city of Machala, El Oro Province, Ecuador (C) the distribution of households surveyed in this study. Household locations were aggregated to the neighborhood level for de-identification. Some clusters (5 households) have been disaggregated across block boundaries. This figure was created in ArcGIS version 10.3.1 (ESRI, 2016) using shapefiles from the GADM database of Global Administrative Areas, version 2.8, freely available at gadm.org. Streets are derived from data available at the OpenStreetMap project (openstreetmap.org) for the municipality of Machala, El Oro, Ecuador. Neighborhood polygons were manually digitized by AMS, and the shapefile data are available upon request to the authors.

### Study design

We conducted diagnostic testing for dengue on all study participants, and we surveyed heads of households regarding household demographics, housing condition, and dengue prevention KAPs. The ascertainment and recruitment of households into this study is described in detail elsewhere [50]. Briefly, individuals who present with clinically-suspected dengue at one of five clinical sites operated by the MOH in Machala, Ecuador are invited to participate in this ongoing study. In this paper, we present an analysis that includes participants from January 2014 through September 2015.

After giving informed consent, the participants were tested for DENV infections using the dengue NS1 rapid strip test (PanBio). A random subset of dengue rapid test-positive individuals (up to four per week) was invited to participate in a household study, and these households are referred to herein as index households. In addition, individuals from the nearest four neighboring households (within 200 meters) in the four cardinal directions were invited to participate in the household study, and are referred to as associate households. A maximum of four individuals per household were invited to participate in the study. This study design was developed and optimized in dengue surveillance studies in Thailand [51,52].

The household study consisted of three parts: a survey of the head of the household, interviewer observation of household characteristics, and blood draw of each household member who was available and who consented for dengue testing. The survey was completed by the head of the household (self-identified), or a proxy (adult age 18 years or greater who was at home during the study team visit, usually the husband or wife) if not available. The survey included questions about the demographics of the head of the household, household demographics, access to water and sewage services, water storage and use practices, knowledge and attitudes about dengue, and prevention activities employed by members of the household. The interviewers’ observations included condition of the house and patio, construction materials, presence and condition of screens in windows and doors, and presence of uncovered standing water on the property. The survey instrument was originally developed in a study that we conducted on household risk factors associated with *Ae. aegypti* [41], and was modified to address risk factors that emerged in a qualitative study of community perceptions [25]. Both of these prior studies were conducted in Machala in 2010-2011. The instrument was developed in Spanish, and the current version was field tested by the study team in households in Machala prior to the start of the study. The survey instrument has been provided in English (Supplementary Text 1) and Spanish (Supplementary Text 2)

Blood samples from study participants were tested for DENV using RT-PCR, NS1 rapid strip test, and PanBio commericial ELISA assays for NS1 and IgM assays. See Stewart Ibarra, 2017, for details of the diagnostic testing procedures [50]. A participant was categorized as having an acute or recent DENV infection if he or she tested positive for any of these tests. Households were characterized as having a DENV infection present if anyone in the household tested positive for DENV. All index households, by definition, had DENV infections present.

### Statistical analysis

Statistical analyses were conducted using SAS version 9.4. Bivariate analyses of households with versus without acute or recent DENV infections were conducted using Chi-square, Fisher’s Exact, or t-tests. Multivariate logistic regression was conducted in two steps using the proc logistic command with backward selection. In the first step, all potential main effects were included in the analysis. In the second step, two-way interactions between all of the variables identified in the first step were added to the analysis.

## RESULTS

We conducted a household-level study to identify KAP and social-ecological risk factors associated with acute or recent DENV infections in the city of Machala, Ecuador. From January 2014 through September 2015, 72 cases of acute DENV infections (NS1 positive) were identified in our surveillance system. A random subset of 44 of these cases (44/72, 61%), along with four neighboring households, were selected to participate in this investigation. Thus, a total of 219 households were included in the household study: 44 index households and 175 associate households (Figure 1). These households were distributed across the city of Machala, thereby representing a range of social-ecological conditions. Most of the households (n=161) were recruited during 2014, and the rest (n=58) were from 2015. The head of the household was female in 24.2% of households, and had a mean age of 47.8 (SD=13.6) years. The households were classified as having (n=139) or not having (n=80) a member with an acute or recent DENV infection. Approximately one third of the households with DENV infections (44/139) were index households.

All of the index households, by definition, had at least one individual who tested positive for DENV. The number of individuals with DENV infections per index household ranged from one to four (mean=1.6). This accounted for 64% of household members with DENV infections on average (range 25-100%). Among the associate households, a range of zero to four (mean=0.58) individuals per household had DENV infections, accounting for a mean of 30% of household members (range 0-100%).

We compared the social-ecological characteristics and reported barriers to dengue prevention in households with versus without DENV infections (Table 1). In bivariate analyses, the presence of DENV infections was positively associated with heads of households who were male, employed, and of younger age than households without dengue (p<0.05). Households with DENV infections were more likely to have a patio with more than 50% shade or to have adjacent abandoned property, and were less likely to have piped water inside of the house or have their trash picked up daily (p<0.05).

**Table 1:**
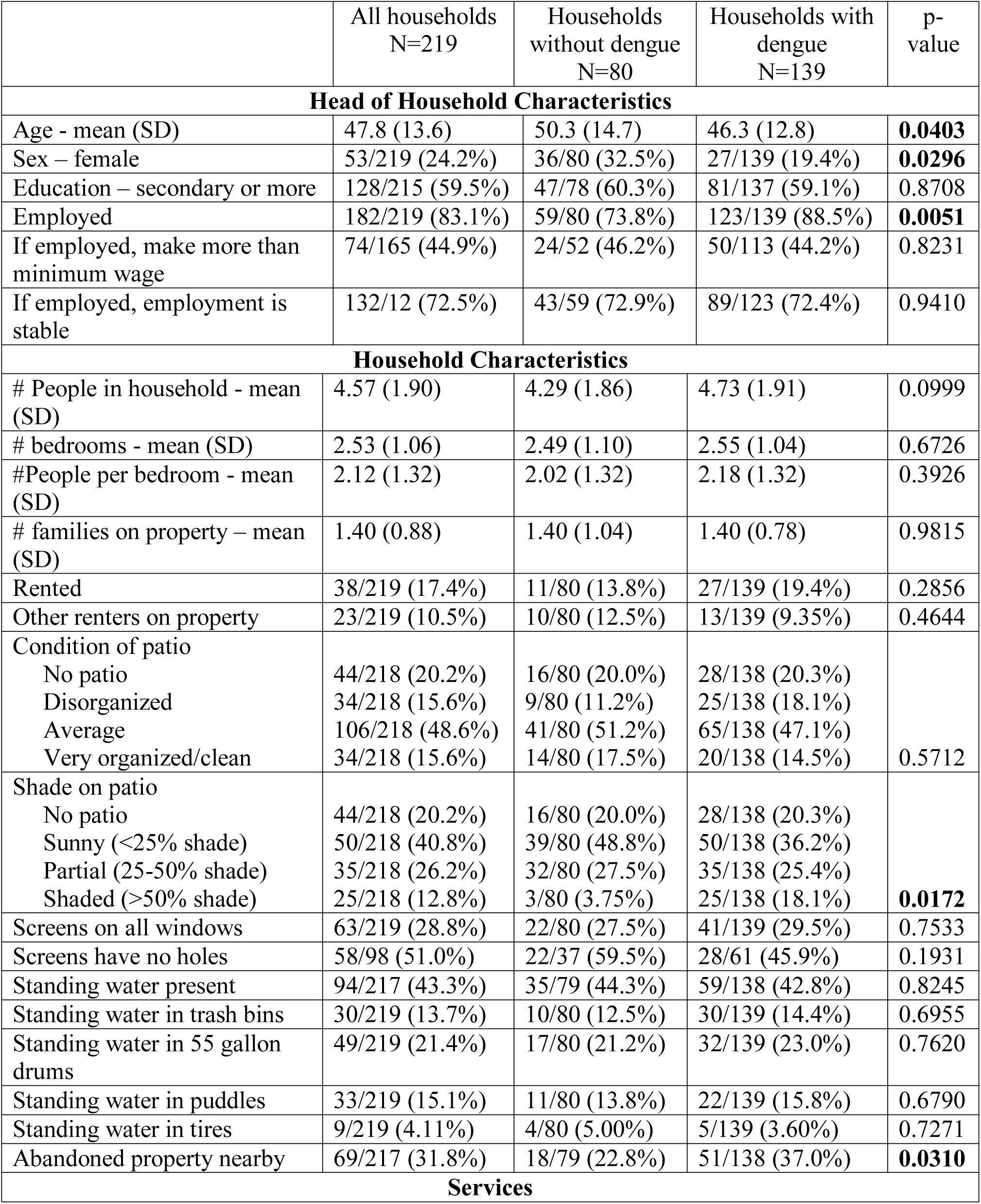
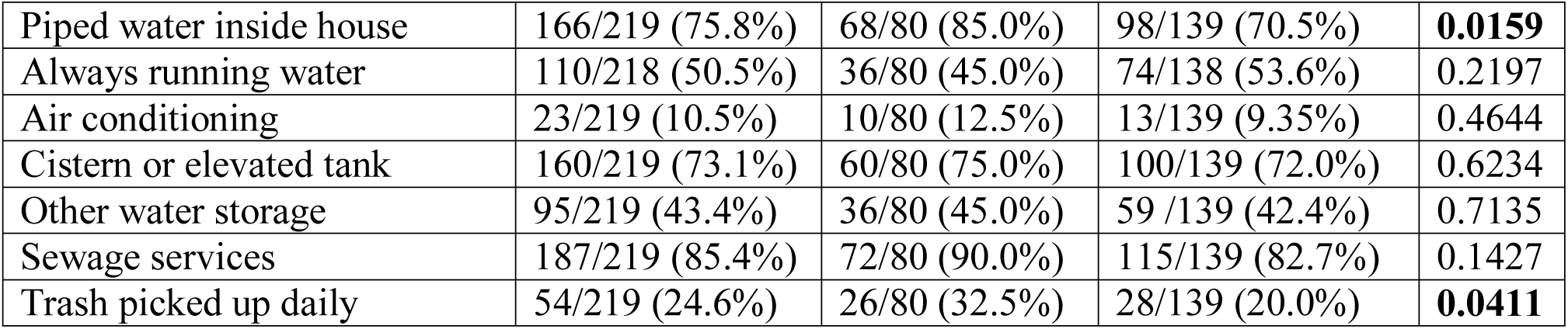
Social-ecological factors in households with versus without acute or recent dengue infections.

|  | All households<br>N=219 | Households<br>without dengue<br>N=80 | Households with<br>dengue<br>N=139 | p-value |
| --- | --- | --- | --- | --- |
| **Head of Household Characteristics** |  |  |  |  |
| Age - mean (SD) | 47.8 (13.6) | 50.3 (14.7) | 46.3 (12.8) | **0.0403** |
| Sex – female | 53/219 (24.2%) | 36/80 (32.5%) | 27/139 (19.4%) | **0.0296** |
| Education – secondary or more | 128/215 (59.5%) | 47/78 (60.3%) | 81/137 (59.1%) | 0.8708 |
| Employed | 182/219 (83.1%) | 59/80 (73.8%) | 123/139 (88.5%) | **0.0051** |
| If employed, make more than minimum wage | 74/165 (44.9%) | 24/52 (46.2%) | 50/113 (44.2%) | 0.8231 |
| If employed, employment is stable | 132/12 (72.5%) | 43/59 (72.9%) | 89/123 (72.4%) | 0.9410 |
| **Household Characteristics** |  |  |  |  |
| # People in household - mean (SD) | 4.57 (1.90) | 4.29 (1.86) | 4.73 (1.91) | 0.0999 |
| # bedrooms - mean (SD) | 2.53 (1.06) | 2.49 (1.10) | 2.55 (1.04) | 0.6726 |
| #People per bedroom - mean (SD) | 2.12 (1.32) | 2.02 (1.32) | 2.18 (1.32) | 0.3926 |
| # families on property – mean (SD) | 1.40 (0.88) | 1.40 (1.04) | 1.40 (0.78) | 0.9815 |
| Rented | 38/219 (17.4%) | 11/80 (13.8%) | 27/139 (19.4%) | 0.2856 |
| Other renters on property | 23/219 (10.5%) | 10/80 (12.5%) | 13/139 (9.35%) | 0.4644 |
| Condition of patio |  |  |  |  |
| No patio | 44/218 (20.2%) | 16/80 (20.0%) | 28/138 (20.3%) |  |
| Disorganized | 34/218 (15.6%) | 9/80 (11.2%) | 25/138 (18.1%) |  |
| Average | 106/218 (48.6%) | 41/80 (51.2%) | 65/138 (47.1%) |  |
| Very organized/clean | 34/218 (15.6%) | 14/80 (17.5%) | 20/138 (14.5%) | 0.5712 |
| Shade on patio |  |  |  |  |
| No patio | 44/218 (20.2%) | 16/80 (20.0%) | 28/138 (20.3%) |  |
| Sunny (<25% shade) | 50/218 (40.8%) | 39/80 (48.8%) | 50/138 (36.2%) |  |
| Partial (25-50% shade) | 35/218 (26.2%) | 32/80 (27.5%) | 35/138 (25.4%) |  |
| Shaded (>50% shade) | 25/218 (12.8%) | 3/80 (3.75%) | 25/138 (18.1%) | **0.0172** |
| Screens on all windows | 63/219 (28.8%) | 22/80 (27.5%) | 41/139 (29.5%) | 0.7533 |
| Screens have no holes | 58/98 (51.0%) | 22/37 (59.5%) | 28/61 (45.9%) | 0.1931 |
| Standing water present | 94/217 (43.3%) | 35/79 (44.3%) | 59/138 (42.8%) | 0.8245 |
| Standing water in trash bins | 30/219 (13.7%) | 10/80 (12.5%) | 30/139 (14.4%) | 0.6955 |
| Standing water in 55 gallon drums | 49/219 (21.4%) | 17/80 (21.2%) | 32/139 (23.0%) | 0.7620 |
| Standing water in puddles | 33/219 (15.1%) | 11/80 (13.8%) | 22/139 (15.8%) | 0.6790 |
| Standing water in tires | 9/219 (4.11%) | 4/80 (5.00%) | 5/139 (3.60%) | 0.7271 |
| Abandoned property nearby | 69/217 (31.8%) | 18/79 (22.8%) | 51/138 (37.0%) | **0.0310** |
| **Services** |  |  |  |  |
| Piped water inside house | 166/219 (75.8%) | 68/80 (85.0%) | 98/139 (70.5%) | **0.0159** |
| Always running water | 110/218 (50.5%) | 36/80 (45.0%) | 74/138 (53.6%) | 0.2197 |
| Air conditioning | 23/219 (10.5%) | 10/80 (12.5%) | 13/139 (9.35%) | 0.4644 |
| Cistern or elevated tank | 160/219 (73.1%) | 60/80 (75.0%) | 100/139 (72.0%) | 0.6234 |
| Other water storage | 95/219 (43.4%) | 36/80 (45.0%) | 59 /139 (42.4%) | 0.7135 |
| Sewage services | 187/219 (85.4%) | 72/80 (90.0%) | 115/139 (82.7%) | 0.1427 |
| Trash picked up daily | 54/219 (24.6%) | 26/80 (32.5%) | 28/139 (20.0%) | **0.0411** |

We also compared KAPs in households with versus without DENV infections (Table 2). The households with versus without DENV infections did not differ on any of the five knowledge and attitude questions, or on reported barriers to dengue prevention activities. We asked survey respondents about whether they engaged in twelve different preventive activities. The most commonly-reported dengue prevention activities were eliminating standing water (37.9%), covering water containers (37.9%), fumigating inside the house (37.9%), cleaning the garbage (37.0%), applying chemicals to standing water (i.e., larvicides) (25.6%) and using mosquito bed nets (20.6%). Households with DENV infections were more likely to report that they applied chemicals to standing water, and were less likely to report the use of indoor fumigation (Table 2). The other prevention activities did not differ between households with versus without DENV infections.

**Table 2.**
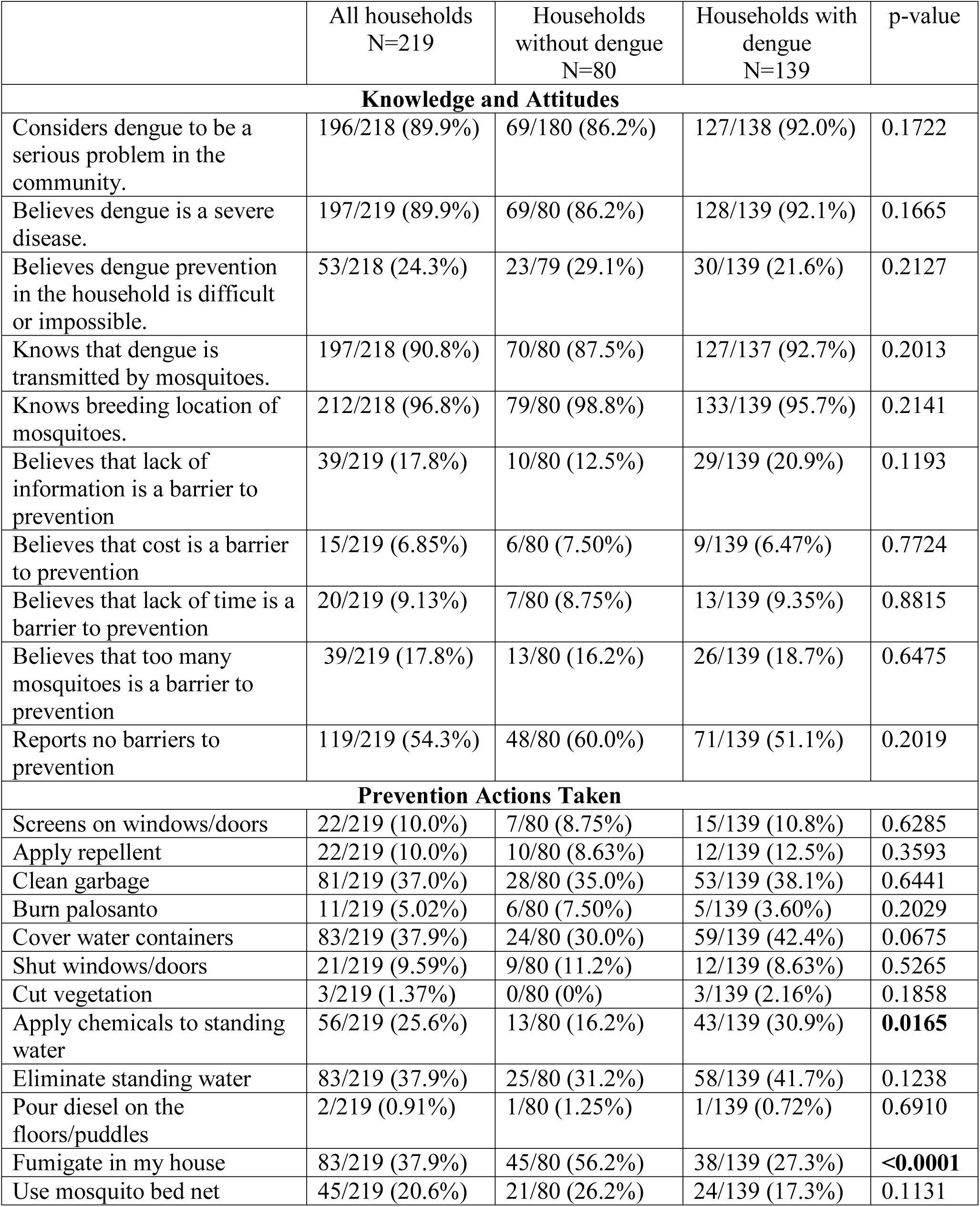
KAPs in households with versus without acute or recent dengue infections. All households Households without dengue N=80

All social-ecological factors and KAPs were used in a logistic regression analysis to identify a multivariate model to predict the presence of an acute or recent DENV infection in the household (Table 3). Model 1 included main effects only, and demonstrated that abandoned properties nearby, frequent interruptions in water supply, and patios with >50% shade were risk factors for dengue. Protective factors in this model included piped water inside the house, fumigation inside the house, use of mosquito bed nets, and reporting cost of as a barrier to protective practices. The strongest factor in this model was the presence of >50% shade on the patio, with an adjusted odds ratio (adj. OR) of 16.2 (95%CI: 2.98-88.1, p=0.001). We then added all two-way interactions of these variables to the model, and eliminated non-significant factors and in a backward selection process. In Model 2, the presence of a patio with more than 50% shade was highly predictive of DENV infections in the household, with an adj. OR of 13.3 (95%CI: 3.2-54.3, p=0.0003), compared to households without a patio or with a patio that had <50% in the shade. Use of mosquito bed nets was protective against DENV infections in this model (adj. OR=0.39. 95%CI: 0.18-0.85, p=0.02). There were two significant interaction terms in Model 2. First, fumigation in the house was protective against DENV infections when there were no abandoned houses nearby (adj. OR=0.19, 95%CI: 0.09-0.42, p<0.0001) but not when there was one or more abandoned properties nearby. Second, fumigation was protective when there was piped water inside the house (adj. OR=0.19, 95%CI: 0.09-0.39, p<0.0001) but not when there was no piped water in the house.

**Table 3.**
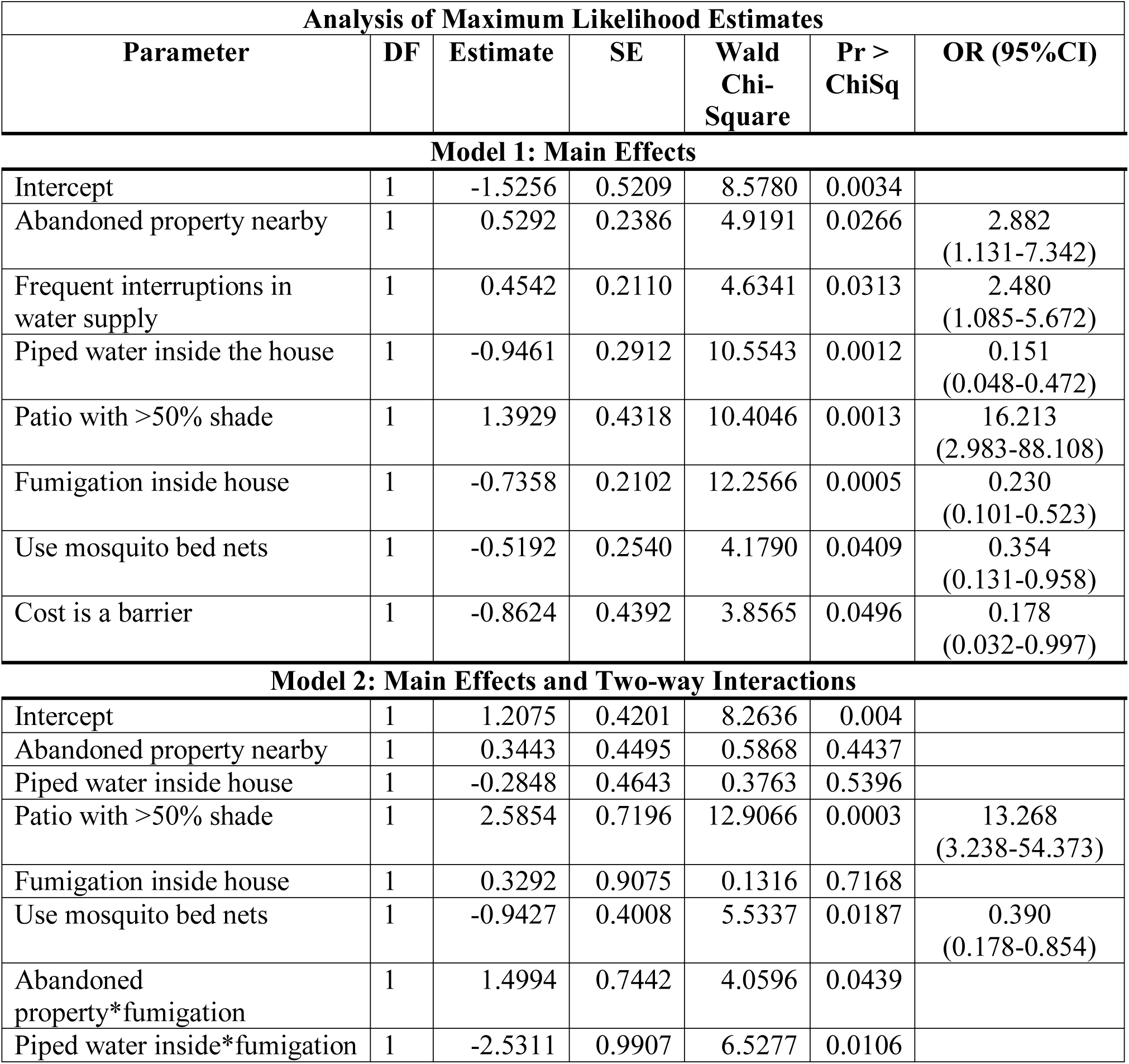
Multivariate logistic regression model of predictors of acute or recent dengue infections in the household.

## DISCUSSION

In this study, we found that specific social-ecological factors and preventive actions were associated with effective dengue control in an region with a high burden of disease, providing important information to guide public health interventions. The strongest predictor of DENV infections in the household, in both multivariate models, was having a patio that was more than 50% shaded. Patio shade and patio condition have been shown to be associated with the presence of *Ae. aegypti* mosquitoes in prior studies, including studies in Machala [41,53]. Our results also support the use of mosquito bed nets, as people who did not report the use of mosquito bed nets were about twice as likely to live in a household that had an acute or recent dengue infection. Other studies have shown associations among dengue infections, KAPs and demographic factors. In our study, with DENV infection as the primary endpoint, demographic variables are not significant factors in the multivariate model. Likewise, knowledge and attitude responses were not associated with DENV infections in the household.

Abandoned properties nearby and lack of piped water inside the house were significant predictors of DENV infections in the house when only main effects were included in Model 1, but not in Model 2. Likewise, fumigation inside the home was found to be protective against DENV infections in Model 1, but only in conjunction with other factors in Model 2. The statistical interactions suggest that the risk factors of abandoned properties and lack of piped water inside the house cannot be overcome with fumigation inside the home. That is, fumigation inside the home is only effective in the absence of abandoned properties nearby and the presence of piped water inside the house. Prior studies in Peru [16] and Australia [17] demonstrated the impacts of indoor residual spraying on a reduction on *Ae. aegypti* densities and dengue risk. In a recent review of indoor residual spraying (IRS) and indoor space spraying (ISS), the authors found that there is evidence of a reduction in *Ae. aegypti* densities, but there is limited evidence of the effects on DENV infections [54]. In our study, 37.9% of participants reported fumigation inside the house as a preventive action, but we did not distinguish between fumigation by the MOH versus by the household members themselves. Many people in Ecuador fumigate their own homes, and there are a variety of products available on the market [55]. It should be noted that a high degree of insecticide resistance in *Ae. aegypti* has been reported in this region (A. Stewart, *pers. comm*.). Resistance is a major public health concern, since insecticides are one of the primary means of controlling *Ae. aegypti* transmitted diseases [56]. Studies are ongoing to document the prevalence of resistance to specific groups of insecticides, to inform vector control interventions.

In this study, 20.6% of households reported the use of mosquito bed nets, and use of nets was associated with a 2.6-fold decreased risk of DENV infections in the household. The protective role of bed nets against DENV infections has been debated in the literature. Studies from rural Thailand reported results similar to this study [57]. Insecticide-treated bed nets offer protection both as a physical barrier during daytime sleeping, and by killing *Ae. aegypti* that come into contact with them, as shown in trials in Haiti [58]. Other studies have failed to find an association between mosquito net use and DENV infections [59,60], presumably because *Ae. aegypti* feeds during the day (morning and afternoon) [61]. However, bed nets are recommended for children napping during the day and to prevent further spread of the virus by viremic individuals resting at home under nets during the day [60,62]. We did not distinguish between insecticide-treated and untreated mosquito bed nets, nor did we gather information on the use of nets (i.e., hours per day, time of day). Based on our local experience, a high proportion of families in the urban periphery use untreated bed nets to protect against nuisance mosquitoes in the early evening and at night. There is a high level of acceptability of the use of bed nets by community members as a result of intensive malaria prevention campaigns in the 2000s. Dengue prevention campaigns in Machala in recent years have not focused on the use of bed nets. However, during the recent epidemic of Zika fever, the MOH targeted the distribution of bed nets in coastal Ecuador to pregnant women. Further research is needed to elucidate the association between mosquito bed nets and DENV protection observed in this study.

There were two additional social-ecological factors that were significant in bivariate analyses but not multivariate model: daily trash pick-up, and application of chemicals to standing water. Applying chemicals to standing water positively correlated with having DENV infections in the household, probably because those who responded “yes” had standing water to begin with. In our experience, chemical application refers to the use of granular organophosphate larvicides (temefos/abate) provided by the MOH and the use of bleach by households to purify the water. Daily trash pick-up appears to be protective against DENV infections in the household, and could be considered by communities as part of dengue prevention programs, even though it was not included in the multivariate model.

In contrast to previous studies in Ecuador, we found that piped water in the house was protective against dengue. Prior entomological field studies and neighborhood-level geospatial analyses of MOH dengue cases in Machala found that access to piped water and poor housing conditions interacted to increase dengue risk [41,42]. When these studies were conducted in 2010-2011, we observed that households that had recently received piped water continued to store water due to poor quality of access and established water storage behaviors. A number of factors could contribute to different findings in the current study. First, the prior studies utilized MOH dengue cases and vector indices as a proxy for DENV risk, which may have introduced biases. Second, it is possible that the quality of piped water (e.g., frequency of interruptions, sediment in the water) improved from 2010 to 2015 due to the major urban renovation projects that occurred during that time. Qualitative improvements in piped water access would reduce the need to store water, thus increasing the protective role of piped water. Third, community members may have changed their water storage behaviors in response to MOH education or other factors.

We also found that DENV infections in the household were associated with younger, male heads of households who were employed outside of the home (in the bivariate analyses). This is in contrast to prior geospatial analyses of MOH dengue cases, which found that neighborhoods with a higher proportion of older, female heads of household were at greater risk [42]. The active surveillance methods in the current study allowed us to more accurately characterize the burden of disease by identifying asymptomatic cases of DENV infection and individuals with DENV infection who had not sought medical care, which were not accounted for in previous studies. Demographic differences between symptomatic and asymptomatic cases may have introduced bias in earlier studies. In Machala, community members reported that working men in the urban periphery are the group least likely to seek healthcare [25], and health care providers [63] supported this notion; therefore, prior studies based on MOH case reports would have underestimated their risk of infection. Also, this study focused on data from individual households rather than neighborhood-level data, and may have therefore been better suited to tease out factors related to the spread of DENV between households. The differences between this study and prior studies highlight the complex nature of DENV transmission and vector control programs.

The results of this study contribute to a growing body of knowledge on the role of social-ecological factors and KAPs on the prevalence of DENV infection. Risk factors vary by location and over time, highlighting the importance of local studies to understand disease risk factors and to inform localized interventions. In a recent case-control study in China, Chen and colleagues showed that living in old apartment buildings increased the risk of DENV infection, while knowledge of dengue fever, use of repellent, and cleaning trash/water containers decreased the risk of DENV infections [43]. In studies from Camaroon, India, the Texas-Mexico border, Sudan, and Key West, Florida, USA, the presence of DENV antibodies (IgG, indicative of a past infection) in individuals was associated with lack of knowledge about dengue fever, high household density (more than three people per bedroom), more than two children in the home, water storage, lack of air conditioning, and poor housing conditions [45–49]. In a Malaysian study, the seroprevalence of DENV IgG in school children was positively associated with apartment/condominium homes and households in a rural setting, while neighborhood fogging, preventive actions and knowledge were associated with the absence of seropositivity in the community’s school children [44]. In a community-level study in Singapore, investigators compared the attributes of communities that were and were not transmission dengue hotspots [64]. They found that protection factors included male heads of households, higher education, having landed property, knowledge of preventive practices, and practicing certain preventive activities (changing water in vases or bowls on alternate days, and removing water from flower pot plates on alternate days).

The main strength of this study is that through a combined passive and active surveillance study design, we focused on laboratory-confirmed acute and recent DENV infections as the primary endpoint. Many prior KAP studies focused on the use of preventive activities, MOH case reports, or vector densities as proxies for dengue risk. We were thus able to include capture and classify asymptomatic infections, as well as symptomatic infections that were not reported to the MOH due to demographic differences in healthcare seeking behavior, factors which may have introduced bias into other studies. In addition, we made use of direct observation in order to capture characteristics of the houses, which eliminates possible errors introduced by self-report. We were also able to triangulate findings from this study to findings from prior qualitative and quantitative studies of dengue risk factors from the same city, allowing us to highlight differences and similarities across the studies.

The main limitation to this study is that we have no way of knowing where the individuals were infected with DENV. In addition to the home, individuals could also have been exposed at other locations such as school or work, and we do not account for risk factors at these locations. In the bivariate analyses, employment by the head of the household was a risk factor for DENV infections in the household, suggesting that exposure at work may be an important factor. A second limitation is that most of the DENV-positive households in this study were index cases, all of whom were referred through MOH health care facilities. Bias related to health care-seeking behavior may have been introduced as a result. Ideally, the analysis would include only associate households, but the sample size would have been too small for statistical analysis. We chose to include all households in the analysis in order to maximize the power of our analysis. A third limitation is that it is not possible to rule out the possibility that members of households with acute or recent DENV infections have recently changed their behavior or risk perception in response to the DENV infection. Biases could have been introduced by self-report or proxy report [65], although as noted earlier, some of the important variables in this study (e.g., shading of patio) were obtained by research staff observation.

Our results suggest that specific actions at the household and community levels could reduce the spread of DENV infections. In resource-limited communities such as Machala, public health actions by the MOH could focus vector control interventions in high-risk households and communities. Tun-Lin and colleagues have shown that targeted interventions based on either types of water containers [66] or conditions of the household [67] can be at least as effective as non-targeted interventions. As in those studies, we found that homes next to abandoned properties and homes with heavily-shaded patios should be high-priority homes for interventions. However, we did not find any specific container type to be associated with DENV infections in the household.

KAP studies have several limitations, such as cultural influences on validity of results [65], and the inability of this tool to capture the complexity of underlying social-political structural drivers that influence DENV infections [68,69]. Factors beyond the individual and community levels play important roles in determining the efficacy of vector control programs [70]. For this reason, the results from KAP studies should be triangulated with data from more comprehensive qualitative approaches in order to understand how and why local risk factors affect disease transmission.

Based on our findings, we suggest that future studies, such as randomized trials, should investigate the impact of the following interventions on acute/recent DENV infections: targeting of vector control in highly-shaded properties, fumigating inside the home, and the use of mosquito bed nets. Community-level interventions include clean-up of abandoned properties, daily trash pick-up, and reliable piped water inside houses. Our results suggest that these community actions moderate the effectiveness of fumigation in prevention of DENV infection, and thus represent very important components of a a community-based approach to prevention. These interventions will require strong inter-institutional collaborations across community leadership councils (responsible for social mobilization), municipal government (responsible for garbage collection, piped water, and abandoned properties), and the MOH (responsible for bed net distribution, fumigation, and other vector control) [25,71]. Our findings also highlight the importance of framing dengue prevention interventions in the context of broader urban development goals (e.g., improve access to piped water), which prior studies showed to be of greater interest to community members [25]. These community- and household-level interventions should also provide some protection against other *Ae. aegypti*-borne diseases such as chikungunya and Zika fever.

## Acknowledgements

This project was possible thanks to support from colleagues from the Ministry of Health, the National Secretary of Higher Education, Science, Technology, and Innovation (SENESCYT) of Ecuador and community members from Machala, Ecuador. We thank our local field team and collaborators for their dedication and perseverance: Jefferson Adrian, Victor Arteaga, Jose Cueva, Reagan Deming, Carlos Enriquez, Prissila Fernandez, Froilan Heras, Naveed Heydari, Jesse Krisher, Lyndsay Krisher, Elizabeth McMahon, Eunice Ordoñez, Tania Ordoñez, and Mercy Silva. Thank you to Clinical Research Management (CRM) for supporting continued surveillance activities. Map data copyrighted OpenStreetMap contributors and available from https://www.openstreetmap.org

## Supporting Information Legends

Supporting Text 1: Household survey instrument in English

Supporting Text 2: Household survey instrument in Spanish

